# Effects of hydroelectric dam construction on spatio-temporal changes of land use land cover in Bui National Park, Ghana

**DOI:** 10.1101/2021.01.28.428667

**Authors:** Godfred Bempah, Prince Boama, Changhu Lu

**Author notes:** Corresponding author;, Current address; Nanjing Forestry University, 159 Longpan Road, Nanjing, Jiangsu, China.

## Abstract

The construction of hydroelectric dams in forest reserves has become a matter of concern for biodiversity conservationists. Visibly among which is the potential to cause changes in climate and land cover and subsequently affect fauna and flora composition. Spatio-temporal changes of climate and land cover in the Bui National Park was analyzed using indices calculations of the landscape based on land cover maps obtained from Landsat satellite images for pre-and post-dam construction periods. Significant changes in land cover following the dam construction were observed. Notable changes include the built-up areas and water body, which recorded an increase of 315.64 % and 4593.43 % respectively, while the forest area decreased. Significant reduction in rainfall (U = 24, *ρ* < 0.05) and increase in temperature (U = 22.5, *ρ* < 0.05) were observed between the pre-and post-dam construction periods. Increased human activities such as illegal mining, indiscriminate tree felling, uncontrolled cattle grazing and charcoal production within the reserve results from inadequate monitoring and law enforcement after the dam construction could likely compound the changes in land cover.

## Introduction

Land use relates to human use of an area of land, that is, whether for settlement, agriculture, forest, wildlife reserve etc. while land cover defines the ecological condition and physical appearance of the land surface (Geist and Lambin 2002) e.g. grassland, forest, water, built-up/bare etc. on an area of land. Both land use and land cover (referred to as LULC) have a common causal agency and changes to the land, affecting both its use and cover. It exerts extensive environmental complications resulting from their combined effects on soil and water quality, biodiversity and microclimate (Schneider and Pontius 2001). Changes in land use types often result in land cover changes; and human activities dominate the land use types, which therefore are the most influential causes of land cover changes (Turner and Meyer 1991).

Other factors such as advancement in technology, wealth creation and changes in economic policies of governments may act in synergy with human population growth to influence LULC changes (Lambin et al. 2003) and this is now receiving considerable attention (Abdelali 2018). One example is the construction of hydroelectric dams as part of the several initiatives to provide solution to the high demand by people for electricity in most countries (Kaunda et al. 2012). Globally, about 900,000 dams have produced about 16.3% of the world’s total electricity supply (IEA 2012) by modifying about 50% of large rivers (Nilsson et al. 2005).

Some negative environmental consequences come with the construction of hydroelectric dams (Beck et al. 2012) including the flooding of large areas which results in loss of biodiversity and riparian ecosystems (Cunha and Ferreira 2012; Woldemichael et al. 2012) and other related global environmental changes (Lambin et al. 2003) as well as increased human activities around dams once the dams start operation (Woldemichael et al. 2012).

Ghana has witnessed great changes in LULC within the past few decades as natural forests and savannah woodlands are converted to different forms such as agricultural lands, built up areas and open land caused by human activities (Antwi et al. 2014) and climate change (Dakwa 2018). At Ghana’s Bui National Park (BNP), the impoundment of the Black Volta River for hydroelectric development is envisaged to cause a major LULC change, involving about 21 percent of land mass of BNP that will be permanently inundated when it is fully impounded and this will lead to excessive effects of biodiversity loss, terrestrial and aquatic ecosystems destruction, and most likely the influx of people into the area to influence land use changes (ERM 2007). The main objective for this study is to assess the spatial-temporal effects of hydroelectric dam construction on the Bui natural ecosystems in order to review and implement environmental policies to manage, conserve and protect resources in the Bui National Park and its environs.

## Method

### Study area

The Bui National Park (80 23’ 13.2072’’ N, 20 22’ 43.9788’’ W) covers an area of about 1,821 km² and it is located close to the Ghana-Cote D’Ivoire border (Fig. 1). It is the third largest wildlife conservation area in Ghana. The Black Volta River bisects the park into two, with a portion lying in the north-western corner of the Bono Region and the other extending towards the southwestern part of the Savanna Region. The park was gazetted in 1971 to primarily provide protection for the drainage basin of the Black Volta River as well as to conserve biodiversity in the entire designated area. The Park is renowned for its high biological richness. The vegetation of the area is mostly Guinea Savannah woodland with patches of Moist Semi-Deciduous Forest, harboring several animal species such as the hippopotamus (*Hippopotamus amphibius*), buffalo (*Syncerus caffer*), and roan antelope (*Hippotragus equinus*). The area experiences a relative humidity of about 75% and yearly rainfall of 1,140 mm (Appiah et al. 2017). The Bui hydroelectric dam is found at the southern end of the BNP. The dam is a gravity, roller compacted concrete type with a height of a 110 m above foundation and creating a reservoir of 12,350 million m² with a total surface area of 440 km² (Habia 2009; Obahoundje et al. 2018).

**Fig. 1.**
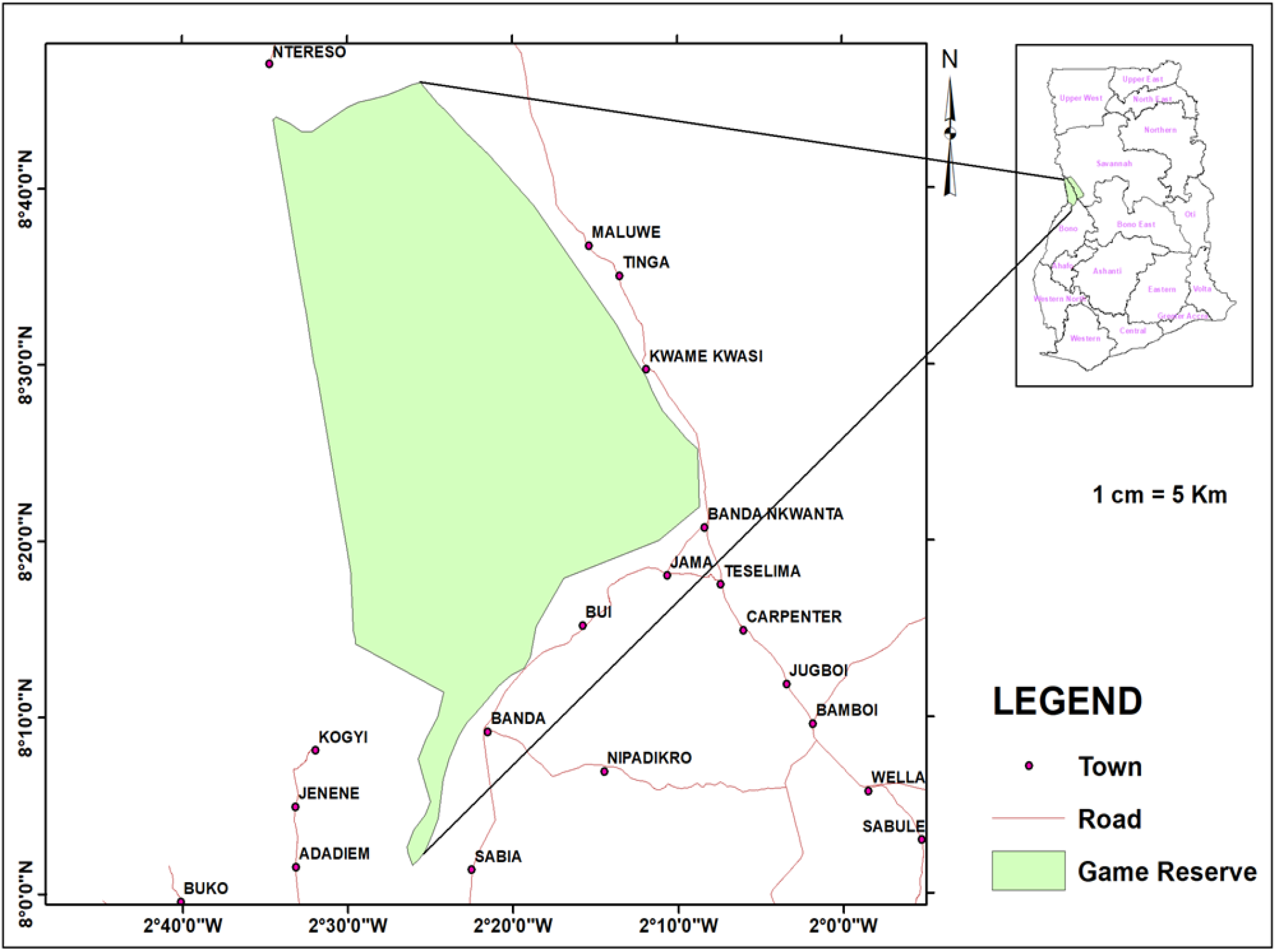
Map of study area

### Data collection

#### Image Acquisition

The land cover maps and matrixes were produced using remote-sensing and Geographic Information System (GIS) application. Landsat 7 Enhance Thematic Mapper Plus (ETM+) and Landsat 8 Operational Land Imager (OLI) images with little or no cloud cover for the years 2000, 2010 and 2020 were obtained from the United States Geological Survey (USGS) Global Visualization Viewer (GLOVIS) website (https://glovis.usgs.gov/). The percentage cloud cover information for each image was obtained from the metadata on the same website. Landsat image was used for this study because of its long coverage of the earth surface (open source and free). All the shape files such as those of the community, road and the park had the same coordinate system, that is, Universal Transverse Mercator (UTM) Zone 30 North. This was to avoid geometric errors associated with working with data with different coordinate systems and ensure consistency and precision. All the satellite images were co-registered to ensure they were aligned with corresponding pixels representing the same objects.

#### Image Preprocessing

Data were prepared to compensate for systematic errors before analyses. Duggin and Robinov (1990) expect such errors to come from sensor spectral properties, atmospheric scattering, among others and to creep into the data acquisition process to reduce the quality of the remote sensor data. Since the errors could cause difficulties in comparing more than one images of the same scene, picked under different conditions, it was important to remove these effects. Therefore, (1) scan lines error associated with Landsat 7 caused by malfunctioning of the sensor were corrected by using the focal analysis tool in Erdas Imagine 2018 to fill in the gaps created by these scan lines. (2) Landsat tiles are made up of bands, each defined by the particular range of wavelength within which it captures radiation to produce image (Lillesand et al. 2015).

The layer stack tool in Erdas Imagine 2018 was used to stack the bands for each tile to generate a composite image to obtain more spectral information from the composite image than analyzing the individual bands separately (Lillesand et al. 2015). (3) Haze has an additive effect to the overall image, resulting in higher digital number (DN) values, and as such, it is reducing the contrast (Lillesand et al. 2015). Its impact differs per band, highest in the blue range, lowest in the IR range (Lillesand et al. 2015). The “haze correction” tool in Erdas Imagine 2018 was used for this operation. This tool identifies the lowest DN value in each band then subtracts it from the remaining DN values for each band. This operation improves the contrast of the images.

The composite images generated from the layer stack operation were used as inputs for this operation. To improve the visualization of the images, image enhancement technique called histogram equalization was performed. This was done in Erdas Image 2018 using the “histogram equalize” tool. This operation further enhances the contrast and makes easy to detect subtle differences among earth surface features identified in the images. (4) The interest of this study was to analyze the land use changes within the study area and since the Landsat images extend beyond its boundary there was a need to extract areas that fell within it for further analysis. The “clip” tool under the data management toolbox in ArcGIS 10.5 was used to extract areas of the composite images (outputs from the haze correction) that fell within the study area.

#### Image Classification

Classification of image was carried out in two modes, unsupervised and supervised classification. This is performed to establish and earmark real world thematic classes to the image pixels (Lillesand et al. 2015).

#### Unsupervised Classification

In the unsupervised classification method, the computer software categorizes the pixels into common land use types using their spectral characteristics without any training data (Lillesand et al. 2015). In this study the isodata unsupervised classification algorithm in Erdas Imagine 2018 was used to classify each clipped image (output from the clip operation) into 30 – 40 classes. The classes were merged based on the spectral similarities to form 8 – 10 classes. The study area, community and road shape files were overlaid on the outputs to check the distribution of these classes within the study area and for easy navigation for field data collection. A map was produced from the 2020 classified image of the study area, which extended to the community and included the roads. This was used as preliminary map for field data collection, during which pixels picked from the unsupervised classification where traced on the field.

#### Field Data Collection

This was necessary to take coordinates of the various land cover types identified as training datasets from the supervised mode. The data collected were numbered serially and divided into two with one half (even numbers) used for training of the images and the other half (odd numbers) for ground truth accuracy assessment of the classified images. The data collected from the field included the coordinates of the points using GPS receiver, the land use description at that point. All the data were entered into a field sheet and later transferred into a Microsoft Excel Sheet. The field data were grouped according to the land cover types identified.

#### Supervised Classification

Here, the pixel categorization process is supervised by an analyst by identifying clearly, to the computer algorithm, numerical descriptors of the various land cover types existing in a location on the field (Lillesand et al. 2015). The classification was done in Erdas Imagine 2018, by using the polygon tool in training the pixels and introducing subclasses of land use types to reduce the margin of error. The maximum likelihood algorithm was then used in running the classification as it considers the cluster center and also its shape, size and orientation by calculating a statistical distance based on the mean values and covariance matrix of the clusters (Bakker et al. 2004). After the classification, the outputs were displayed in ArcGis 10.5 and the subclasses were together using the reclass tool. The reclassified maps were then filtered using the “majority filter” tool in ArcGIS 10.3.

#### Accuracy Assessment and Change Detection

The field data were the reference data used to assess the accuracy of the classification and the confusion matrix and kappa statistics used to produce statistical outputs to check the quality of the classification results. Information on variations corresponding to trend, spatial distribution and extent of change can be obtained from change detection applications (Singh 1989). The “matrix union” tool in Erdas Imagine 2018 was used to detect pixel by pixel changes to the final land cover maps. Thus it picked a pixel from the old land use map and compared it to the same pixel on the new land use map and so change matrix were generated from the change maps, which provided information on change detection in hectares of land cover or use.

#### Rainfall and Temperature data

Climatic variables such as temperature and rainfall for the BNP are monitored by the Bole-Bamboi District Meteorological Office. Monthly records on temperature and rainfall from 2000 to 2020 used for this study were obtained from the District Meteorological Office.

#### Data analysis

Data on climate (Rainfall and temperature) and land cover changes over a period between 2000-2020 were analyzed using time series statistics with the Paleontological statistics software package for education and data analysis, PAST version 3.0 (Hammer et al. 2001). Mann-Whitney U test and Kruskal - Wallis tests were performed to determine whether there was a statistically significant difference among them. Observation of lagged correlations among climatic variables time series and land cover changes time series was done using the cross-correlation function (CCF).

## Results

### Rainfall and temperature changes over the period

The highest decade mean temperature (27.69 ± 0 .219 °C) was recorded for 2011–2020 compared to the decade 2000–2010 with 27.16 ± 0.108 °C. The highest decade mean rainfall (93.96 ± 3.094 mm) was recorded for 2000–2010 compared to 2011–2020 with 85.954 ± 2.77 mm. Results of Mann Whitney U test showed that the differences in the two decade were significant for mean rainfall (U = 24, *ρ* < 0.05) and mean temperatures (U = 22.5, *ρ* < 0.05). There was a negative correlation (r = −0.609, Fig. 2), and estimated value from the series’ cross-correlation function (CCF) was significant (*p* = 0.003) between the mean rainfall time series and the mean temperature time series for the study period.

**Fig. 2.**
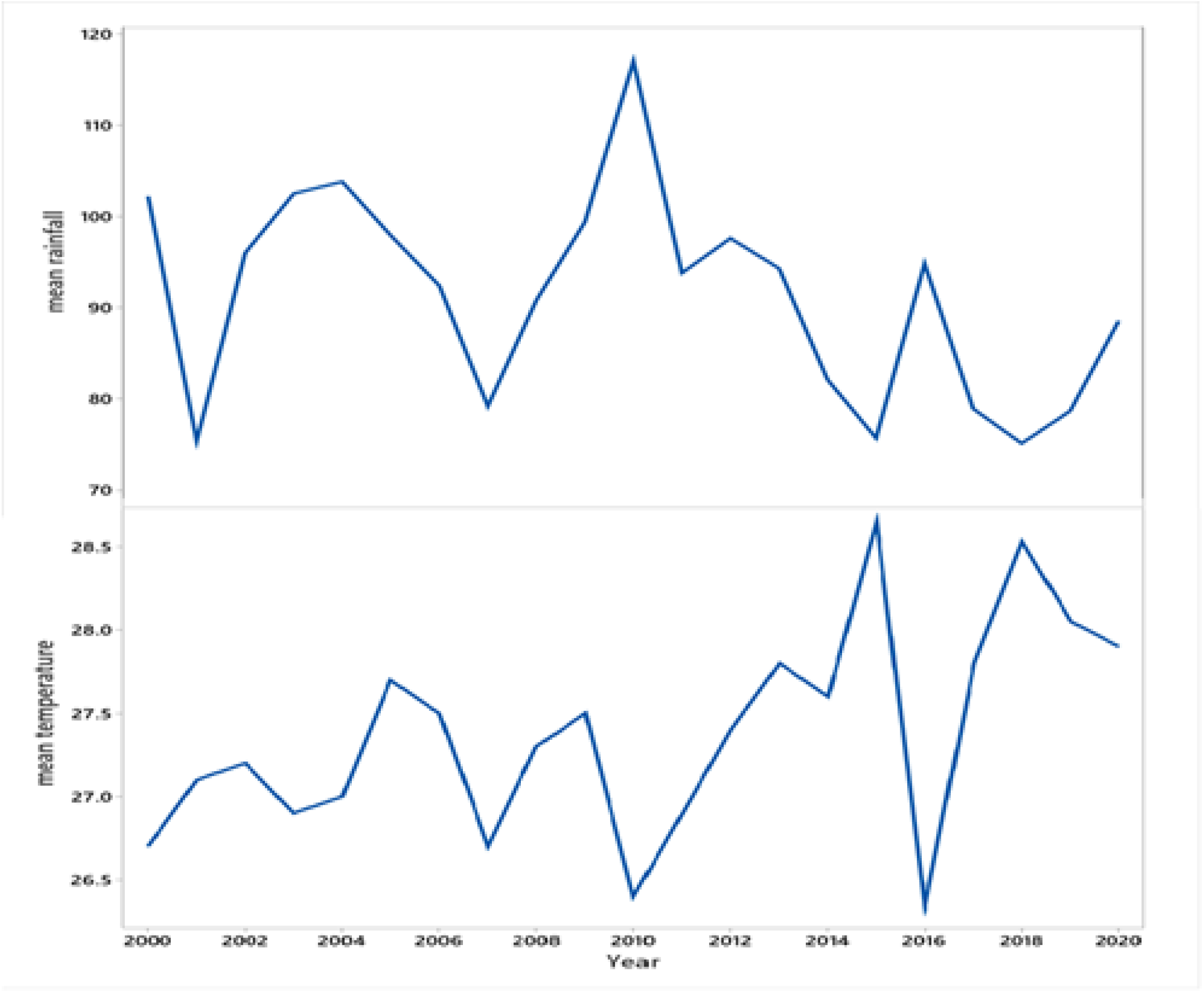
Trends of changes of mean rainfall (mm) and mean temperature (°C) from 2000 – 2020 at BNP

### Land cover changes

In all, five land cover types were identified, namely; built-up, close forest, grassland, open forest, and water body. A set of three different land cover maps covering the study area for the years 2000, 2010 and 2020 is shown in Figure 3. Kruskal Wallis test indicated significant difference among the various landcover classes (H = 9.33, *ρ* < 0.05) and Tukey’s *post hoc* analysis revealed a significant difference between five out of the 10 pairs of the land cover classes. Before the dam construction, open forest occupied the largest portion of the total land area with 45.87 % (Fig. 4). However this changed, during the post-dam construction, with grassland (60.54 %) occupying the largest percentage cover, followed by water body with 19.01 %.

**Fig. 3.**
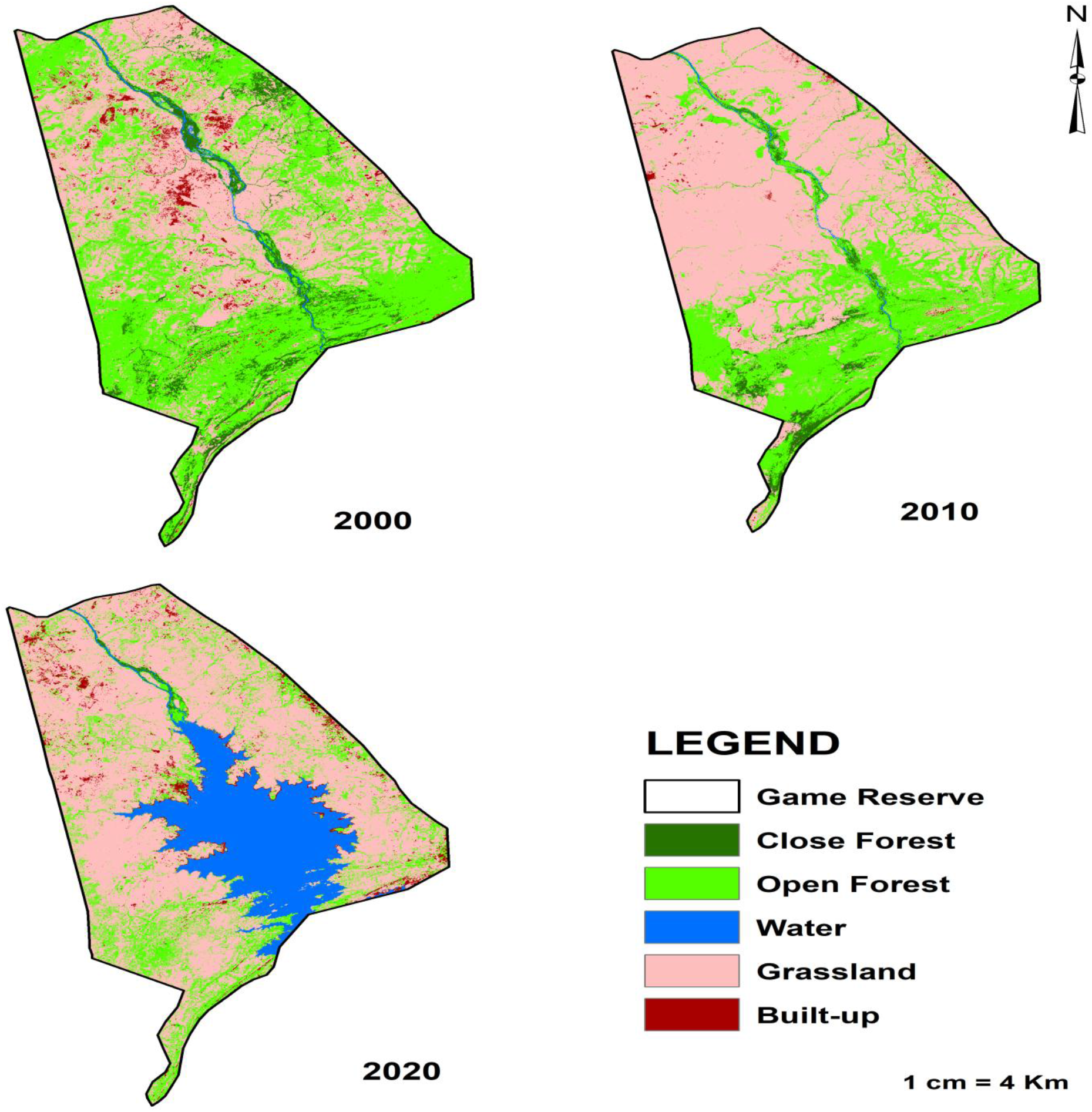
Land cover maps of the Bui National Park for the years 2000, 2010 and 2020

**Fig. 4.**
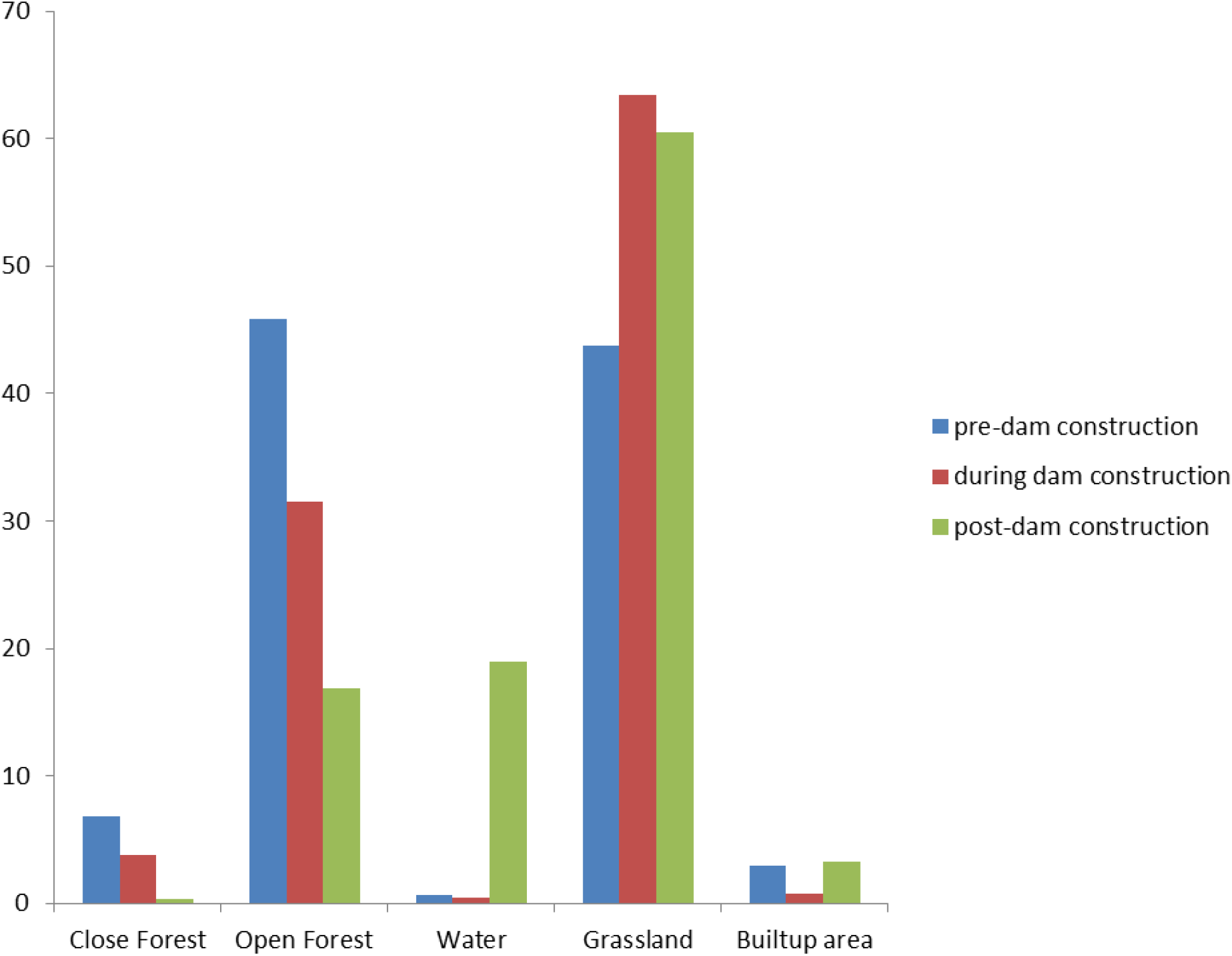
Percentage of area covered by each Land cover types; pre, during and post dam construction

About 38.53 % of the total land cover remained stable while about 61.47 % was converted during the study period. The highest area change was the water body, which increased by 4,593.43 %, from 752.69 ha in 2010 (during dam construction) to 35,326.39 ha in 2020 (post-dam, Table 1). Close and open forest areas decreased from 12,647.01 ha and 85,243.78 ha in the year 2000 to 575.73 ha and 31,282.72 ha in 2020 respectively, representing 95.45 % and 63.3 % decreases respectively (Table 1). During the same period, grassland increased with the variation ratios reaching 38.32 %. Noticeably, built-up areas decreased by 72.55 % before the dam construction but significantly increased by 315.08 % after the construction of the dam (Table 1).

**Table 1:**
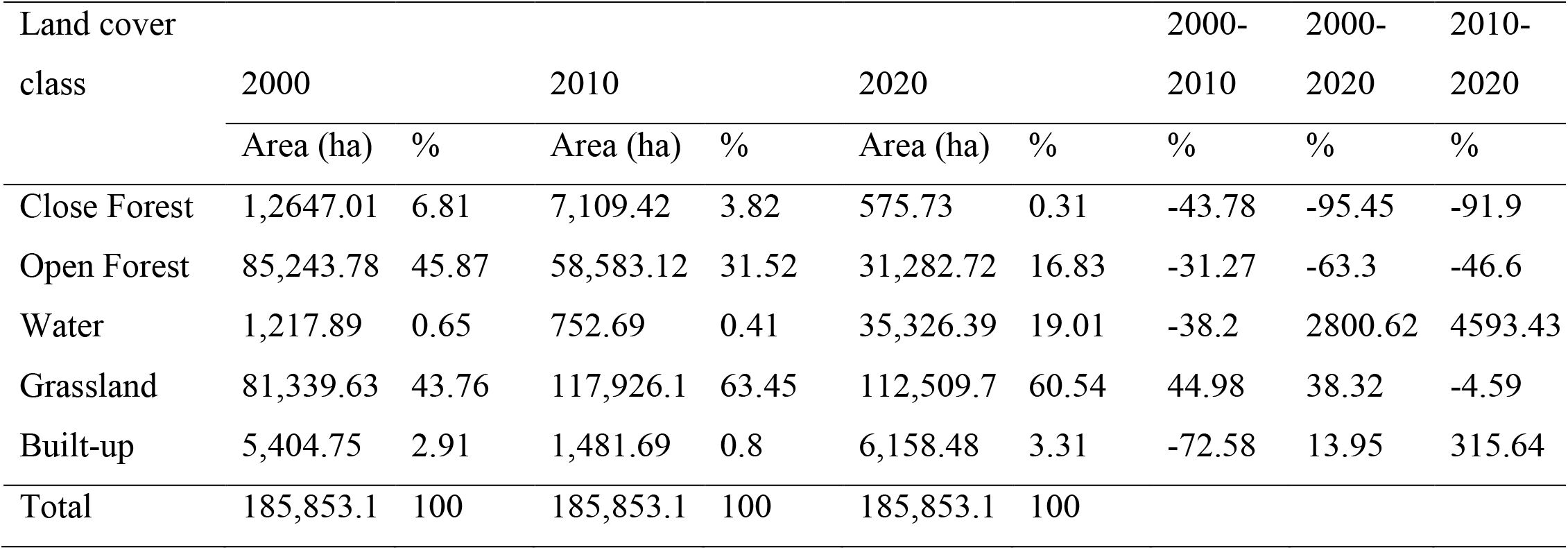
Area and percentage changes of land cover in different year (ha, %).

### Climatic variables and land cover time series over the dam construction periods

Land cover changes and climatic variables seem to follow a similar trend over the period after the dam construction (Fig. 5). For example, a decrease in the amount of rainfall corresponded with a decrease in close forest and open forest. During the same period, water body, grassland and builtup area increased with marginal increase in temperature. Increase in water body results from the reservoir created by the dam and grassland and builtup area is caused by forest degradation corresponding with temperature increase.

**Fig. 5.**
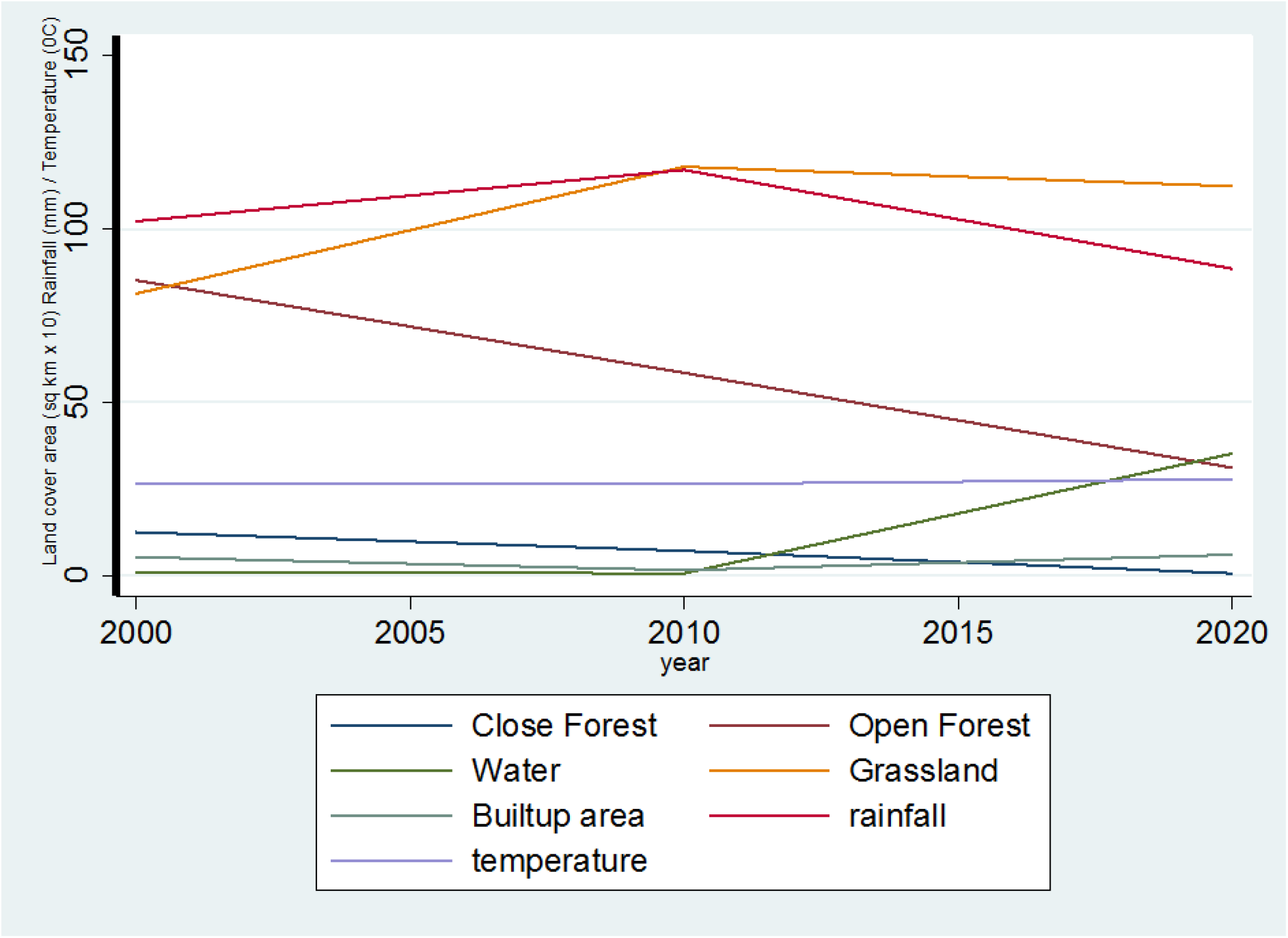
Land cover, rainfall and temperature changes with the construction of the dam

### Accuracy assessment

The overall accuracy for 2000, 2010 and 2020 were 86.67 %, 81.67 % and 83.33 % respectively (Table 2). The highest validation data for 2000, while that of 2010 were of least quality was recorded. The figures indicate that the land cover classification for the periods 2000, 2010 and 2020 were correctly classified. The calculated Kappa coefficients were 81 %, 73.89 %, and 77.32 %, respectively, for 2000, 2010 and 2020 (Table 2), representing strong agreement.

**Table 2:** Accuracy assessment results for 2000, 2010 and 2020

| Land cover | Year |  |  |  |  |  |
| --- | --- | --- | --- | --- | --- | --- |
|  | 2000 |  | 2010 |  | 2020 |  |
|  | Producer accuracy | User accuracy | Producer accuracy | User accuracy | Producer accuracy | User accuracy |
| Close Forest | 0.7143 | 0.7143 | 1 | 0.8571 | 1 | 1 |
| Open Forest | 0.9091 | 0.8 | 0.7778 | 0.7 | 0.7692 | 0.625 |
| Water | 1 | 1 | 1 | 1 | 1 | 1 |
| Grassland | 0.85 | 0.9444 | 0.7692 | 0.8333 | 0.76 | 0.8261 |
| Built-up | 0.8333 | 1 | 0.8333 | 1 | 0.875 | 1 |
| Overall accuracy | 0.8667 |  | 0.8167 |  | 0.8333 |  |
| Percentage accuracy | 86.67 |  | 81.67 |  | 83.33 |  |
| Kappa coefficient | 0.81 |  | 0.7389 |  | 0.7732 |  |

## Discussion

There were significant changes in climate and land cover of the Bui National Park (BNP) over the study period, from the year 2000 to 2020. The decrease in forest cover (close and open) corresponds to decrease in rainfall after the dam construction Also, marginal increase in temperature corresponds with increase in grassland and builtup areas. Dam construction at BNP caused decrease in forest cover, which is consistent with the national rate of deforestation occurring in Ghana (Antwi et al. 2014) and with the results of other studies (Appiah et al. 2017; Zhao et al. 2010) that found a decrease in forest cover after dam construction. Contrarily, Wiejaczka et al. (2017) found no significant changes in forest cover during pre-and post-dam construction.

Several factors may account for the deforestation rate in BNP including; (1) ineffective monitoring of contractors who were permitted to fell trees in the flood zones and therefore allowing indiscriminate felling of trees in the national park; (2) laxity in enforcing laws that protect national park most especially during and after construction of the dam; (3) lack of collaboration and cooperation among institutions responsible for managing the area and stakeholders, to team up to formulate integrated plans to secure the area during and after the project.

Post-dam (2010-2020) reduction in forest cover was more noticeable than pre-dam (2000-2010) reduction. Tree felling activities started from 2008 suggesting that between 2008 and 2010, tree felling activities were officially on going. It is then possible to explain that the deforestation that occurred before 2010 were largely due to preparation towards the construction of the dam. Tree felling activities were still carried out even after the completion of the dam, possibly resulting in the significant decrease in forest cover. Except for the trees found at the riverbanks, it is highly impossible to find a cluster of economic tree species in the study area. This explains the severity of the situation in terms of biodiversity conservation as well as climate change.

These migrants cut trees to build their houses and also as source of energy (firewood) for cooking. Trees were also cut to put up accommodation for resettled communities as well as for staff of the Bui National Park and Bui hydroelectric dam (Appiah et al. 2017). A significant number of illegal miners settled in the area, as well as cattle herdsmen, fisherfolks and charcoal producers. Many small villages are scattered inside the reserve, and several others exist along the fringes. The huge number of people present in and outside the BNP carrying out several activities (Appiah et al. 2017) is linked to exploitation of forest resources on a daily basis (Sharma et al. 2019). These anthropogenic activities are escalating, affecting the reserve resources due to the continuous illegal entry of people into the reserve.

The water bodies increased significantly during the post-dam construction period, which can be attributed to the increase in the level of the reservoir created by the hydroelectric dam in the BNP, an observation that has been reported by other similar studies elsewhere (Obahoundje et al. 2018; Wiejaczka et al. 2017; Zhao et al. 2010).

The grassland decreased its cover from 2010-2020. Other researches (Chalise et al. 2019; Wiejaczka et al. 2017; Xu and Chi 2019; Özüpekçe 2019) have reported similar occurrence of post-dam grassland decline in protected areas. Illegal mining, uncontrolled cattle grazing and charcoal production could account for the decline of grassland during this period. Periodic movement of thousands of cattle within the reserve occurs and this leads to grass trampling and overgrazing. This causes reduction in quality and quantity of forage as well as modifying vegetation and soil formation due to soil compaction (Obahoundje et al. 2018; Panthi et al. 2017; Sharma et al. 2019). In some cases, cattle herdsmen attempt to cause regeneration of fresh pasture by consciously setting fire to the vegetation resulting in uncontrolled destruction of vegetation (Sharma et al. 2019).

The changes in the built-up areas were significant during post-dam construction period, and consistent with reports of several studies (Zhao et al. 2010; Özüpekçe 2019). The increase in built-up areas can be attributed to the emergency of illegal mining observed during field data collection. In nearly all cases of the reports as well as this study, gold mining is cited as the cause of increased sizes of the built-up areas and the incidence of illegal mining is attracting many young people to the area. A serious issue of encroachment lingers on, in the study area contributing to occurrence of built-up/bare lands.

## Conclusion

The post-dam changes resulted in land use and climate changes at the national park. Notable of this are decrease in rainfall, increase in temperature, reduced size of forests, increased water covered surface and increased human settlements. The changes involved conversion of one land cover type to another leading to transfer of sizeable cover from one type to another to cause a decrease or increase of any particular type involved. The activities of the human settlements in particular are likely to compound the changes in landcover negatively. Therefore, a greater cooperation between the BNP and Bui Power Authority, which are the two institutions managing the area, is suggested, to ensure committed enforcement of protective laws and prevent further migration of people into the protected area.

## Acknowledgement

We would like to thank the management and staff of Bui National Park for the assistance provided to us during the field data collection.

## Data Availability Statement

The data presented in this study were obtained from the United States Geological Survey (USGS) Global Visualization Viewer (GLOVIS) at https://glovis.usgs.gov/

## Author contributions

All authors contributed to the study conception and design. Material preparation, data collection and analysis were performed by Godfred Bempah, Prince Boama and Changu Lu. The first draft of the manuscript was written by Godfred Bempah and all authors commented on previous versions of the manuscript. All authors read and approved the final manuscript.

